# A distinct population of L6 neurons in mouse V1 mediate cross-callosal communication

**DOI:** 10.1101/778019

**Authors:** Yajie Liang, Wenzhi Sun, Rongwen Lu, Ming Chen, Na Ji

## Abstract

Through the corpus callosum, interhemispheric communication is mediated by callosal projection (CP) neurons. Using retrograde labeling, we identified a population of layer 6 (L6) excitatory neurons as the main conveyer of transcallosal information in the monocular zone of the mouse primary visual cortex (V1). Distinct from L6 corticothalamic (CT) population, V1 L6 CP neurons contribute to an extensive reciprocal network across multiple sensory cortices over two hemispheres. Receiving both local and long-range cortical inputs, they encode orientation, direction, and receptive field information, while are also highly spontaneous active. The spontaneous activity of L6 CP neurons exhibits complex relationships with brain states and stimulus presentation, distinct from the spontaneous activity patterns of the CT population. The anatomical and functional properties of these L6 CP neurons enable them to broadcast visual and nonvisual information across two hemispheres, and thus play a major role in regulating and coordinating brain-wide activity events.

## Introduction

As the largest bundle of axonal fibers in the mammalian brain, the corpus callosum mediates interhemispheric communications through axonal projections between cortices of the frontal, parietal, occipital, and temporal lobes^1^. Early anatomical studies suggested that callosal projection (CP) neurons are primarily L2/3 or L5 neurons projecting homotopically to the contralateral cortex^2, 3, 4^. Later functional studies indicated that these neurons participate in collaborative processing of information across the hemispheres. For sensory cortices, the transcallosal pathways are thought to enable the processing of bilateral sensory stimuli. For example, CP neurons in the somatosensory cortex were found to mediate bilateral integration of tactile information^5, 6^. In the auditory cortex, CP neurons were shown to contribute to sound localization in spatial hearing^7^.

In visual cortices, callosal connections highly concentrate at the borders between the primary and secondary visual cortices^2, 8, 9^. Because this border region contains a representation of the vertical midline of visual field, these CP neurons, mainly located in L2/3 and L5, were considered to be involved in binocularity and fusion of visual fields^8, 10, 11, 12^. Inactivation experiments indicated that these callosal inputs strongly modulate visual cortical responses^12, 13, 14^. However, whether and how the monocular V1 contributes to callosal projection remains little known. In rodents, the monocular part of V1 was initially considered as acallosal^2^. CP neurons were later found throughout the deep infragranular layer in V1^15, 16^. Compared to other cortical cell types, very little is known about the identity, connectivity, or activity of these CP cells^17^.

In this study, we used a high-efficiency recombinant AAV variant to gain genetic access to CP neurons in the mouse monocular V1, studied their connectivity profiles with a combination of viral strategies, and characterized their activity in awake mice using *in vivo* calcium imaging. We found that V1 CP neurons were concentrated in L6 and formed a distinct population from the *NTSR1*-positive corticothalamic (CT) L6 neurons. Instead of being homotopic, L6 CP neurons formed an extensive network, projecting to and receiving inputs from multiple cortical regions of different sensory modalities. We used rabies viral tracing to identify their presynaptic partners and found cells in both local V1 circuit and long-range cortical areas. Although a substantial proportion of L6 CP neurons encoded visual features such as orientation tuning and possessed well-defined receptive fields, an even larger fraction exhibited spontaneous activity that was often modulated by the presence of visual stimuli. Whereas the spontaneous activity of CT population in the dark was highly positively correlated with the arousal level of the animal, we found that the spontaneous activity of CP neurons exhibited a richer repertoire, suggestive of a multisensory or higher cognitive origin.

## Results

### CP neurons in mouse V1 are dominantly located in L6

Traditionally, CP neurons were labeled with intra-parenchymal injection of retrograde tracers, such as horseradish peroxidase or Fluoro-Gold (FG), with varying efficacy^18^. To label CP neurons with high efficacy, we took advantage of a recently developed recombinant AAV variant (rAAV2-retro) that permits efficient retrograde access to projection neurons^19^. We injected rAAV2-retro carrying GFP (rAAV2-retro.CAG.GFP) into the monocular zone of right V1 of a transgenic mouse line with L5 excitatory neurons labeled with H2B-mCherry (Rbp4-Cre × Rosa26 LSL CAG H2B mCherry^20^). We observed brightly labeled CP neurons in the left monocular V1 predominantly located in the cortical layer below L5, with few cells in the supragranular layer (L2/3) (**Figure 1A**). More superficial CP neurons were observed at the border of V1 and V2L, consistent with earlier studies^2, 8, 9^. Immunostaining with anti-GABA antibody indicated that these neurons were not GABAergic (**Supplementary Figure 1A and 1B**). Therefore, CP neurons in mouse V1 are dominantly excitatory neurons located in L6.

**Figure 1.**
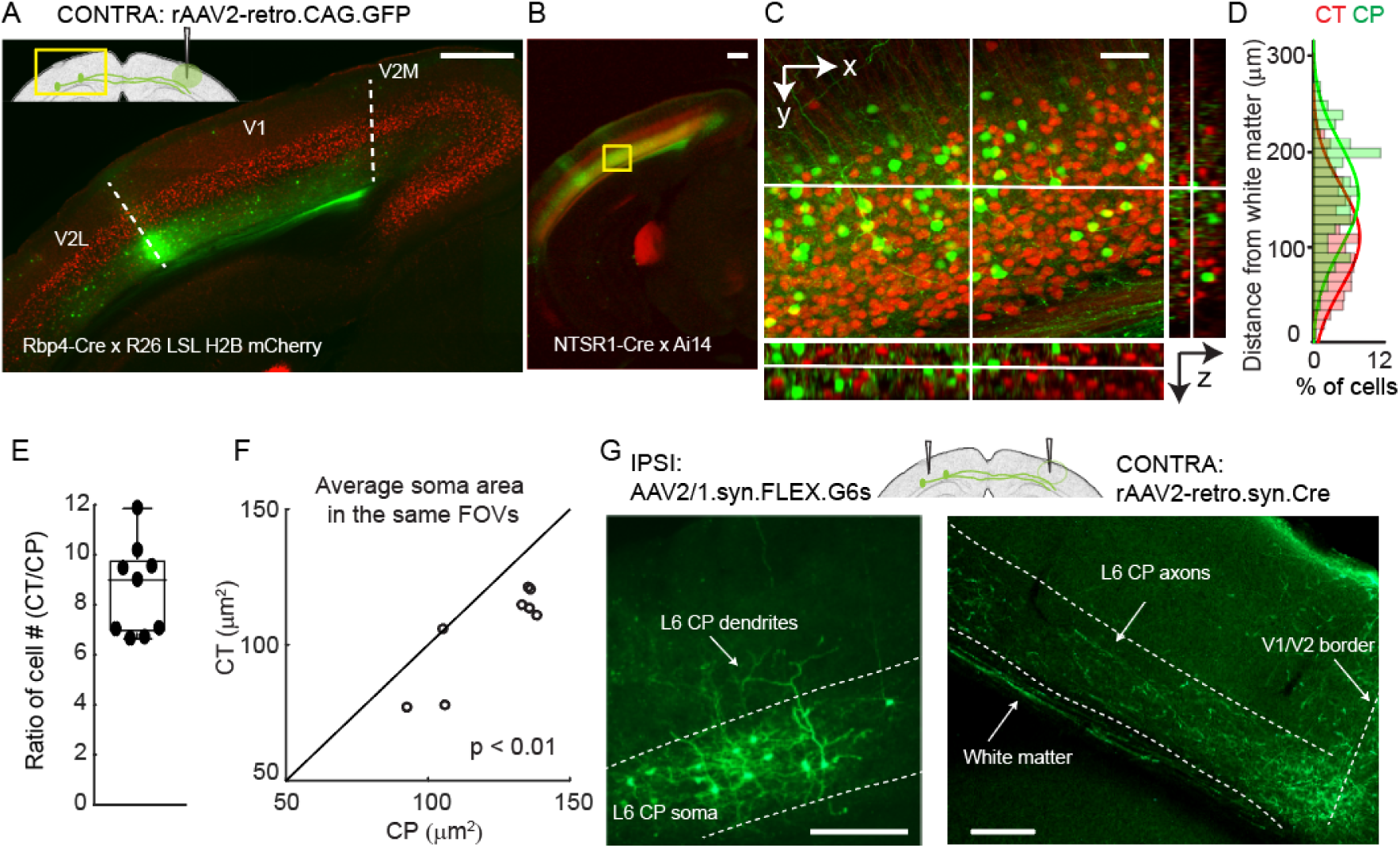
A distinct population of callosal projection (CP) neurons in L6 of the mouse monocular V1. (A, B) Fluorescence images of left V1 after rAAV2-retro.CAG.GFP injection in contralateral (CONTRA) V1 with (A) L5 pyramidal neurons labeled with H2B-mCherry (Rbp4-Cre × R26 LSL H2B mCherry) and (B) L6 corticothalamic (CT) neurons labeled with tdTomato (NTSR1-Cre × Ai14), respectively. (C) Magnified and orthogonal views of the yellow box area in (B) (CP in green, CT in red). (D) Depth-dependence of CT and CP somata distributions measured by cell counting from the volume in (C). 515 CT neurons and 80 CP neurons were counted. (E) Ratio of cell counts (CT versus CP) from the same fields-of-view (FOVs), n = 4 mice (1-2 FOVs from each mouse, 4,718 CT and 611 CP neurons in 9 FOVs). (F) Cell size comparison between CP and CT neurons in the same FOVs. 4,440 CT and 802 CP neurons in 9 FOVs from 4 mice. Wilcoxon signed-rank test, p = 0.0078. (G) Contralateral injection of rAAV2-retro.syn.Cre and ipsilateral injection of AAV2/1.syn.FLEX.GCaMP6s labeled CP neurons (left panel) in ipsilateral (IPSI) V1 and (right panel) their axons in contralateral V1 (axon image taken after additional immunostained with anti-GFP antibody). Dashed lines: (A, G) V1/V2 borders and cortical layers. Scale bar: 500 µm in (A, B); 50 µm in (C); 200 µm in (G).

### L6 CP neurons and *NTSR1*-positive CT neurons are distinct populations

Having identified L6 neurons as the main callosal-projecting neurons in mouse monocular V1, we then asked whether these CP neurons were distinct from the thalamus-projecting L6 CT neurons. To this end, we utilized a Cre-recombinase transgenic mouse line NTSR1-Cre (NTSR1-cre GN220) that selectively labels L6 CT neurons in V1^21, 22, 23^. Crossed with the Cre reporter line Ai14^24^, the resulting mice had L6 CT neurons expressing red fluorescent protein tdTomato. Injecting rAAV2-retro.syn.GFP in the right V1 and labeling CP neurons in the left hemisphere, we similarly observed that CP neurons were mostly located in L6 (**Figure 1B**), with high-resolution image stacks showing a clear separation of CP and CT neurons in V1 (**Figure 1C**). Similar separation was also observed with FG as the retrograde tracer (**Supplementary Figure 1C and 1D**).

Counting the number of cells along the cortical depth, we found slightly different distributions for these two groups of neurons: more CP neurons were found in superficial L6, while there were more CT neurons at depth (**Figure 1D**). In the same fields-of-view (FOVs), CT neurons were ∼nine times denser than CP (median = 9.0, interquartile range IQR = 3.0, 4 mice, 9 FOVs, **Figure 1E**). On average, CP neurons had significantly larger somata (Wilcoxon signed-rank test, p < 0.01, 10 FOVs from 4 mice, **Figure 1F**), consistent with the data from CP neurons in the rat somatosensory cortex^25^. Contralateral injection of rAAV2-retro.syn.Cre and ipsilateral injection of AAV2/1.syn.FLEX.GCaMP6s labeled CP neurons with green fluorescent protein GCaMP6. The dendrites of some CP neurons extended into L5 (**Figure 1G, left panel**) and their projections to the contralateral cortex are mainly localized in the infragranular layers (**Figure 1G, right panel**).

### L6 CP neurons contribute to a horizontal network interconnecting multiple cortical areas across the two hemispheres

Having discovered that V1 receives inputs from L6 CP neurons of contralateral V1, we then asked whether there were other sources of transcallosal inputs into V1. We unilaterally injected rAAV2-retro.CAG.GFP into V1 of NTSR1-Cre × Ai14 mice, where L6 CT neurons expressed tdTomato, and found that in addition to contralateral V1, neurons in secondary visual cortical areas (V2M and V2L), auditory cortex (AuC), and ectorhinal cortex (Ect) in the contralateral hemisphere also projected to V1 (**Figure 2A**). As in contralateral V1 (**Figure 2B**), the CP neurons in the other cortical areas were also distinct from CT neurons (**Figure 2C, 2D**). Interestingly, the CP populations in the contralateral primary and secondary visual cortices were all located in L6 (**Figure 2C**), whereas the CP neurons from AuC and Ect appeared to concentrate in L5 (**Figure 2D**).

**Figure 2.**
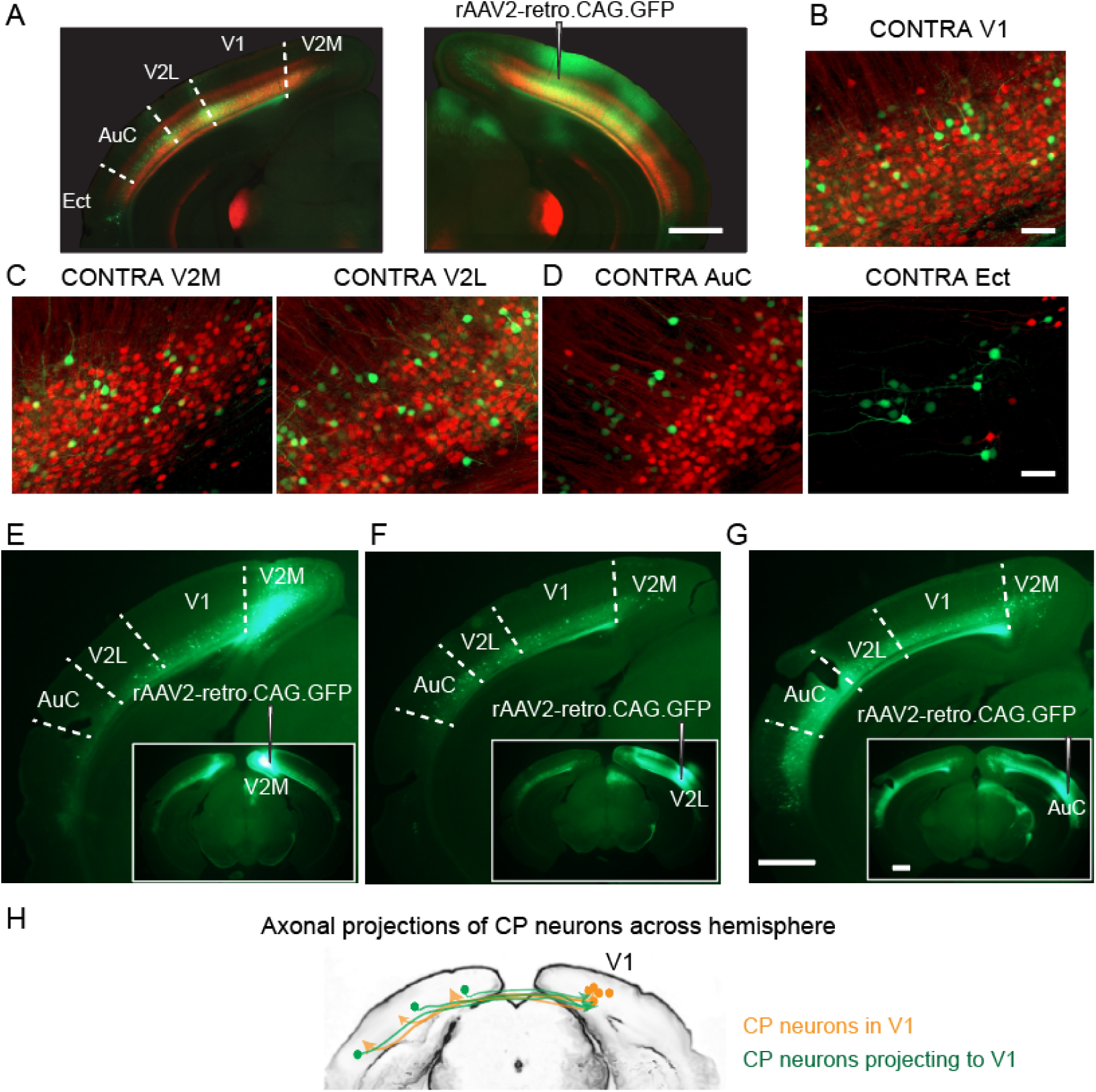
Extensive callosal projections to and from V1. (A) Unilateral injection of rAAV2-retro.CAG.GFP into right V1 of NTSR1-Cre × Ai14 mice, with CT neurons expressing tdTomato (red) and CP neurons in multiple cortical areas of the left hemisphere labeled with GFP (green). (B, C, D) Example coronal images of left V1, V2M, V2L, AuC, and Ect, respectively. n = 2 mice. (E, F, G) Example coronal images of retrogradely labeled L6 CP neurons after injection of rAAV2-retro.CAG.GFP into V2M, V2L, or AuC. n = 2 mice for each group. (H) Schematic showing callosal projection patterns to and from V1. Scale bars: 1 mm in (A, E, F, G); 50 µm in (B, C, D).

Injecting rAAV2-retro.CAG.GFP in V2M, V2L, or AuC cortices, we found similarly widespread staining of CP neurons in the contralateral cortex spanning multiple sensory modalities (**Figure 2E, 2F,** and **2G**). Similar to V1, the extrastriate cortical areas used to be considered as acallosal due to their lack of superficial CP neurons^9^. Here we found that their visual cortical transcallosal communication was similarly subserved by L6 neurons, consistent with another study using a chemical tracer^26^. In AuC, in addition to CP neurons in contralateral AuC, L6 neurons in V1, V2M, and V2L also provided transcallosal inputs. We observed a bias in the CP neuron distribution with respect to sensory modality: more CP neurons were found between cortices processing sensory information of the same type than different types (e.g., visual vs. auditory). Together, these results indicated that V1 not only received widespread transcallosal projections from contralateral cortical regions, but also sent projections to these areas (**Figure 2H**) via L6 neurons. Therefore, L6 CP neurons contribute to a horizontal network interconnecting cortical regions representing multiple sensory modalities across the two hemispheres, and may play a role in interhemispheric cross-modal sensory integration.

### L6 CP neurons receive both local and long-range cortical inputs

The above retrograde labeling experiments with rAAV2-retro vectors revealed the diverse transcallosal projection targets of L6 CP neurons in V1. We next investigated their upstream inputs using rabies viral tracing^27^. We used rAAV2-retro.syn.cre to express Cre recombinase in L6 CP neurons in wildtype mouse V1. We also utilized the Scnn1a-Cre transgenic mouse line that expressed Cre recombinase in L4 pyramidal neurons in V1. Injections of Cre-dependent AAV helper vectors drove expression of glycoprotein (G), avian receptor protein (TVA), and blue fluorescent protein (BFP) in the starter cell population expressing Cre. Three weeks later, we injected a glycoprotein deficient form of the rabies virus encapsulated with the avian sarcoma and leucosis virus envelope protein (ΔRV), which expressed mCherry in neurons presynaptic to the starter cells as well as the starter cells themselves. Consequently, the starter cells were identified as those expressing both BFP and mCherry, and the presynaptic cells were those labeled only with mCherry. The mice were perfused a week later and the brains sectioned and imaged to chart the spatial locations of the presynaptic neurons (**Figure 3A**).

**Figure 3.**
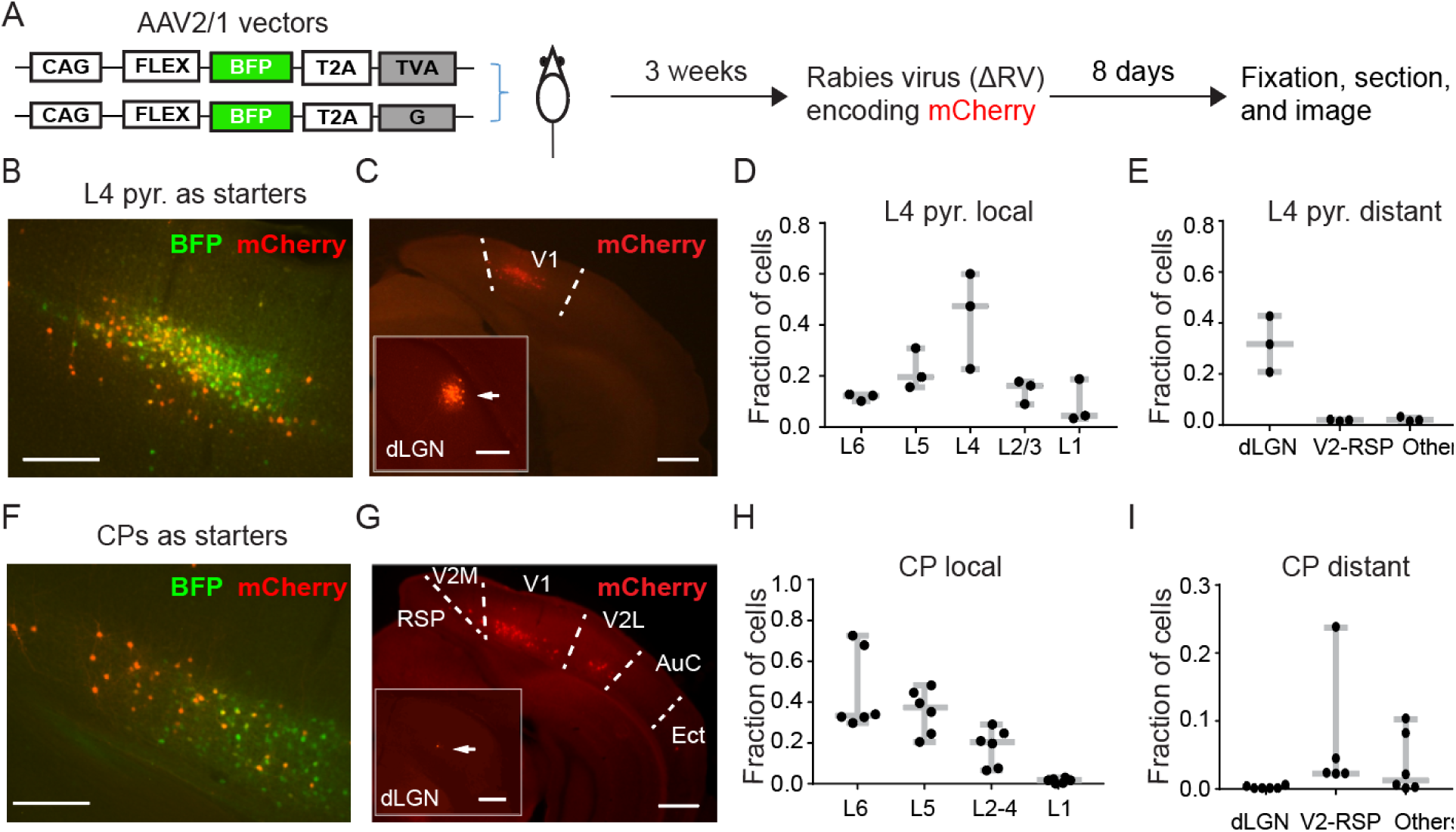
Presynaptic partners of V1 L6 CP neurons from the ipsilateral hemisphere. (A) Schematic showing the experimental procedure. Starter cells were double-labeled by BFP and mCherry. mCherry^+^-only labeling indicated presynaptic partners of starter neurons. (B, C) Example fluorescence images of coronal brain sections with (B) L4 pyramidal neurons as starter cells and (C) their presynaptic neurons ipsilateral to the injection site. Arrows in insets of (C) pointing to presynaptic cells in dLGN. (D, E) Fractions of (D) local (within V1) and (E) distant presynaptic neurons of L4 pyramidal neurons. (F, G) Example fluorescence images of coronal brain sections with (F) CP neurons as starter cells and (G) their presynaptic neurons ipsilateral to the injection site. Arrows in insets of (G) pointing to presynaptic cells in dLGN. (H, I) Fractions of (H) local (within V1) and (I) distant presynaptic neurons of L6 CP neurons. Scale bars: 200 µm in (B, F) and insets of (C, G); 500 µm in (C, G).

To validate our rabies tracing method, we investigated the presynaptic cell population for L4 pyramidal neurons. With L4 pyramidal cells as the starter cells (**Figure 3B**), we observed prominent local (493 out of 726 mCherry^+^ cells, n = 3 mice, **Figure 3C** and **3D**) and long-range presynaptic cells (especially in dLGN, 214 out of 726 mCherry^+^ cells, n = 3 mice, **Figure 3C** inset and **Figure 3E**). This agreed with the known connectivity of these neurons^28^ and confirmed the validity of our rabies tracing method.

With CP cells in V1 as starter cells (**Figure 3F**), we found strong local V1 inputs (2,150 out of 2,459 mCherry^+^ cells, n = 6 mice, **Figure 3G** and **3H**), as well as substantial ipsilateral long-range inputs from V2 (V2M, V2L) and retrosplenial cortex (RSP) (262 out of 2,459 mCherry^+^ cells, n = 6 mice) as well as auditory cortex (AuC) and ectorhinal cortex (Ect) (47 out of 2,459 mCherry^+^ cells, n = 6 mice; **Figure 3G** and **3I**). We also found a few presynaptic neurons in the contralateral hemisphere located in the infragranular layer (**Supplementary Figure 2**). These results indicate that CP neurons receive both local V1 and long-range cortical inputs. Together with the rAAV2-retro experiments, these results suggested that CP neurons receive both local and long-range cortical inputs, while simultaneously transferring information to and integrating information from CP neurons in the contralateral hemisphere. Thereby, they form a functional network that mediates cross-callosal information processing, allowing V1 in each hemisphere to receive information from multiple cortical areas in its own as well as from the contralateral hemispheres.

### Visually-evoked responses of L6 CP and CT neurons in V1 of awake mice

Having characterized the L6 CP neurons anatomically, we then investigated their visually-evoked responses and compared them with L6 CT neurons, using *in vivo* calcium imaging (**Figure 4**). L6 CT neurons in V1 of the left hemisphere were transfected with the calcium indicator GCaMP6s^29^ by injecting AAV2/1.syn.FLEX.G6s in V1 of the NTSR1-Cre mouse (**Figure 4B**). L6 CP neurons were labeled using the same approach as described above (AAV2/1.syn.FLEX.GCaMP6s in left V1 and rAAV2-retro.syn.Cre in the contralateral V1, **Figure 4D**). Presenting drifting gratings to the right eye (100% contrast, 6 s drifting gratings interleaved with 6 s stationary gratings, 0.07 cycles per degree, 2 Hz, 12 directions with 10 trials each in a pseudorandom sequence), we then measured changes in fluorescence brightness (ΔF/F) in the cell bodies of the GCaMP6s^+^ neurons in quietly awake mice after habituating them to head fixation. Imaging CP and CT neurons at depths ranging from 550 to 650 µm below dura with a homebuilt two-photon fluorescence microscope optimized for deep imaging^30^ (**Figure 4C**, **4E**), we identified neurons with detectable calcium transients. A neuron was considered as visually-evoked if its activity during at least one drifting grating stimulus was significantly higher than their activity during the inter-stimulus stationary-grating period by paired t test (p < 0.01)^31^; otherwise, it is considered to be non-visually evoked (spontaneously active). Under this criterion, we found 62% of active CT neurons (792/1282, 10 mice) and 26% of active CP neurons (483/1876, 17 mice) to exhibit visually evoked responses, and focused our current analysis on these visually responsive neurons.

**Figure 4.**
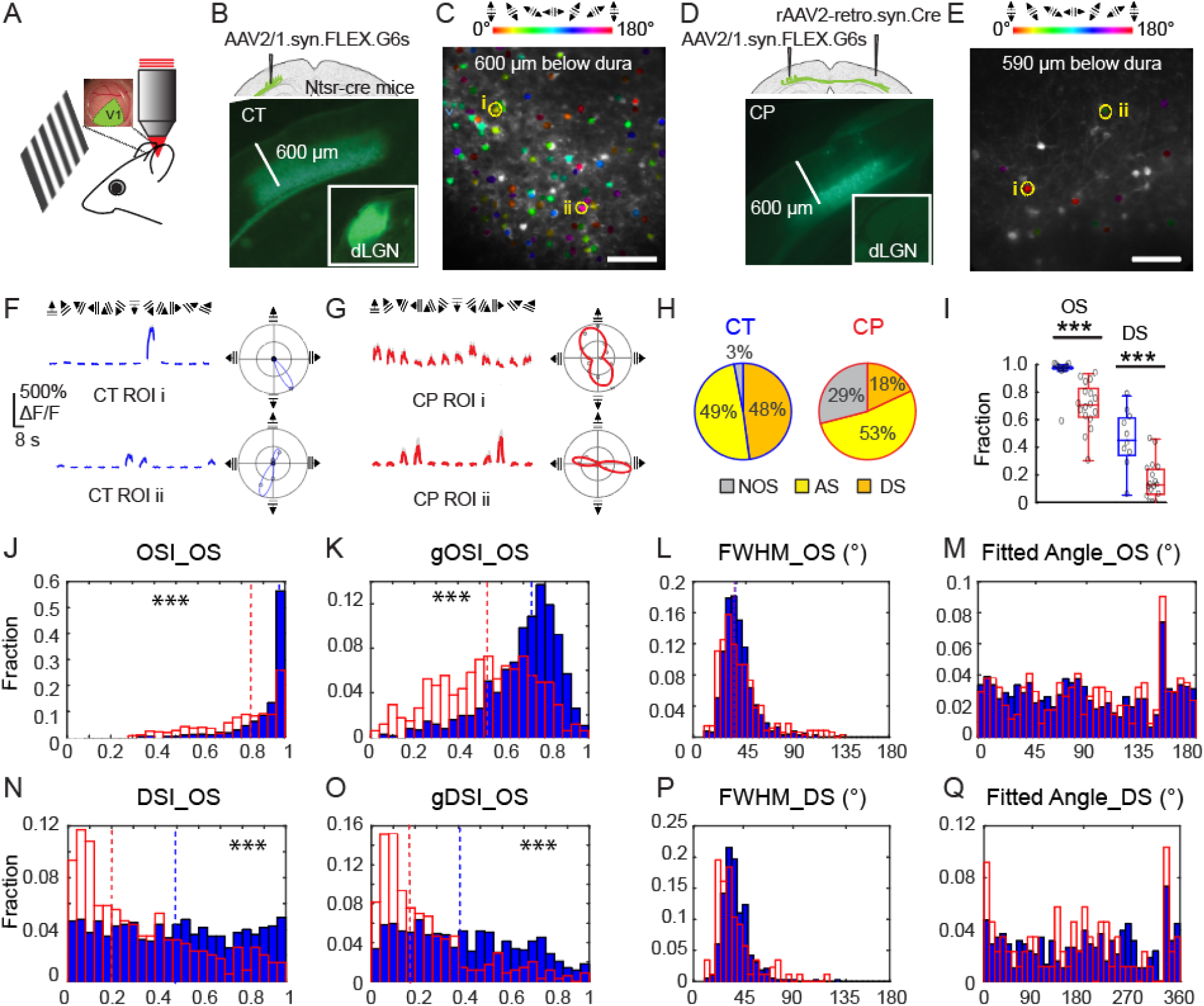
*In vivo* calcium imaging of L6 CT and CP neurons in V1 of the awake mouse. (A) Schematic of *in vivo* imaging and visual stimulation setup. (B, D) Viral labeling strategies and widefield fluorescence images of coronal sections containing GCaMP6s^+^ L6 CT and CP neurons, respectively. Insets: dLGN. (C, E) Example *in vivo* two-photon excitation fluorescence images of L6 CT and CP neurons, respectively, with the orientation-selective (OS) neurons color-coded by their preferred orientation. (F, G) (Left) Example visually evoked calcium responses from two OS CT and CP neurons (i and ii, labeled in C and E), respectively. Colored lines and gray shades: averages and s.d. from 10 trials; (Right) Polar plots of the responses and fitted turning curve for same neurons. (H) Percentages of OS, including axis selective (AS) and direction selective (DS), and non-orientation selective (NOS) CT and CP neurons. CT: n = 796 active visually evoked cells, from 10 mice. CP: n = 483 active visually evoked cells, 17 mice. (I) Fraction of OS and DS cells per mice for CT (blue) and CP (red) neurons. n = 10 mice for CTs; n = 17 mice for CPs. Wilcoxon rank-sum test (nonparametric test): ***p<0.001. (J-Q) Histogram distributions of orientation and direction tuning parameters for CT (blue) and CP (red) neurons, including (J) OSI, (K) gOSI, (L) turning curve FWHMs of OS neurons, (M) preferred orientations of OS neurons, (N, O) DSI and gDSI of OS neurons, (P) FWHM of DS neurons (P), and (Q) preferred directions for DS neurons. Dashed lines: medians. Wilcoxon rank-sum test: ***p<0.001. Scale bars: 600 µm in (B, D); 100 µm in (C, E).

Consistent with an earlier electrophysiological study^32^, we found CT neurons to select for grating orientation and drifting direction (**Figure 4F**). We also found CP neurons whose activity depended on the orientation and moving direction of the drifting gratings (**Figure 4G**). Color-coding the preferred orientation of these orientation-selective (OS) neurons, we found a “salt-and-pepper” pattern in their tuning maps for both CT and CP neurons (**Figure 4C, 4E**), similar to superficial layers of mouse V1^33^. We also explored the relationship between the tuning maps of the two groups of neurons. We labeled CP and CT neurons with GCaMP6s and jRGECO1a^34^, respectively, using Cre- and FLPo-recombinase strategies (**Supplementary Figure 3A, 3B**), and performed calcium imaging on them simultaneously. OS tests did not reveal obvious relationship between the tuning maps of CPs and CTs (**Supplementary Figure 3C-3E**).

Comparing CP and CT neurons with visually evoked responses, we found a smaller proportion of CP neurons to have orientation selectivity: whereas a remarkably high percentage (97%) of CT neurons were OS (out of 796 CT cells from 10 mice), 71% of CPs were OS (out of 483 CP cells from 17 mice) (**Figure 4H**). The same trend held for individual mice, with a significantly higher fraction of OS CT neurons than CP neurons (CT: 0.97 ± 0.03, CP: 0.71 ± 0.22; median ± IQR; 10 CT and 17 CP mice; rank-sum tests, p < 0.001; **Figure 4I**).

We quantified orientation selectivity of each cell with orientation-selectivity index (OSI) and global OSI index (gOSI) (see Methods, **Figure 4J, 4K**). Both OSI and gOSI distributions of OS CT neurons had significantly greater medians than those of CP neurons (CT gOSI: 0.74, OSI: 0.97, 772 neurons from 10 mice; CP gOSI: 0.54, OSI: 0.84, 343 neurons from 17 mice; rank-sum tests, p < 0.001 in both cases). As indicated by the OSI distributions and consistent with the distributions of the full width at half maximum (FWHM) of the tuning curves (**Figure 4L**), both CT and CP populations contained highly orientation-selective and sharply tuned neurons, with their preferred orientations distributed over the entire orientation range (**Figure 4M**). However, we found more broadly tuned CP neurons (FWHM around 100°, e.g., ROI i in **Figure 4G**), which were absent from CT population.

Some OS neurons exhibited substantially different responses towards gratings drifting along opposite directions. Defining neurons with direction selectivity index DSI > 0.5 (or a response ratio towards opposing directions larger than 3, see Methods) as direction-selective (DS), we found about half of the CT neurons to be DS, whereas only a quarter of the orientation-selective CP neurons were DS (**Figure 4H, 4I**). Consistent with this result, DSI and gDSI distributions of OS CP cells had smaller medians than the CT cells (DSI CP: 0.25, CT: 0.49; gDSI CP: 0.17, CT: 0.40; 343 CP neurons from 17 mice, 772 CT neurons from 10 mice; rank-sum tests, p < 0.001 in both cases; **Figure 4N, 4O**). The FWHMs of DS neurons had similar distributions to those of the OS neurons (**Figure 4P**), and their preferred motion directions were distributed throughout the whole range of angles (**Figure 4Q**).

Using sparse-noise stimuli consisting of a pair of white and black squares randomly distributed on a gray background, we also mapped the visual receptive fields of the CT and CP neurons in anesthetized mice, using our previously published method^31^. Cells with defined receptive fields were found in both groups (**Supplementary Figure 4**). For CT neurons, out of 45 cells (n = 3 mice) that showed visually evoked activity, we found well-defined RFs for 27 cells (i.e., 60%); For CP neurons, out of 71 cells (n = 5 mice) that had visually evoked activity, 21 cells were found to have RFs (i.e., 30%).

The existence of L6 CP neurons with visually evoked activity, selectivity towards visual features, and well-defined receptive fields, as revealed by the calcium imaging experiments, suggested that L6 CP neurons in the monocular V1 encode and transmit orientation, direction, as well as receptive field information across the corpus callosum to contralateral cortex. A distinct population from the thalamus-projecting CT neurons, CP neurons nevertheless possess similar orientation tuning characteristics and thus serve as a pathway of visual information flow between V1s of the two hemispheres.

### CP neurons exhibit diverse patterns of spontaneous activity both in the dark and during drifting grating stimuli

While investigating the orientation selectivity of L6 neurons with drifting gratings, we discovered that the calcium transients of 74% of active CP neurons (1393/1876 CP cells in 17 mice) were not synchronized with stimulus presentation, thus appeared to be spontaneously active. In comparison, a much smaller fraction of active CT neurons (38%, 486/1282 CT cells in 10 mice) exhibited such non-visually evoked activity (**Figure 5A**). Keeping the animal in the dark, we found that 83% of CP neurons (1351/1625 neurons from 11 mice) were spontaneously active, a higher proportion than that of CT neurons (48%, 173/359 neurons from 5 mice) (**Figure 5B**). Furthermore, in the dark, the calcium transients of CP neurons have higher amplitudes than CT neurons (mean ΔF/F% during the entire imaging session, median values: CP 14.7, 1351 neurons from 11 mice; CT 6.6, 173 neurons from 5 mice, p < 0.001, Wilcoxon rank sum test; **Figure 5C**).

**Figure 5.**
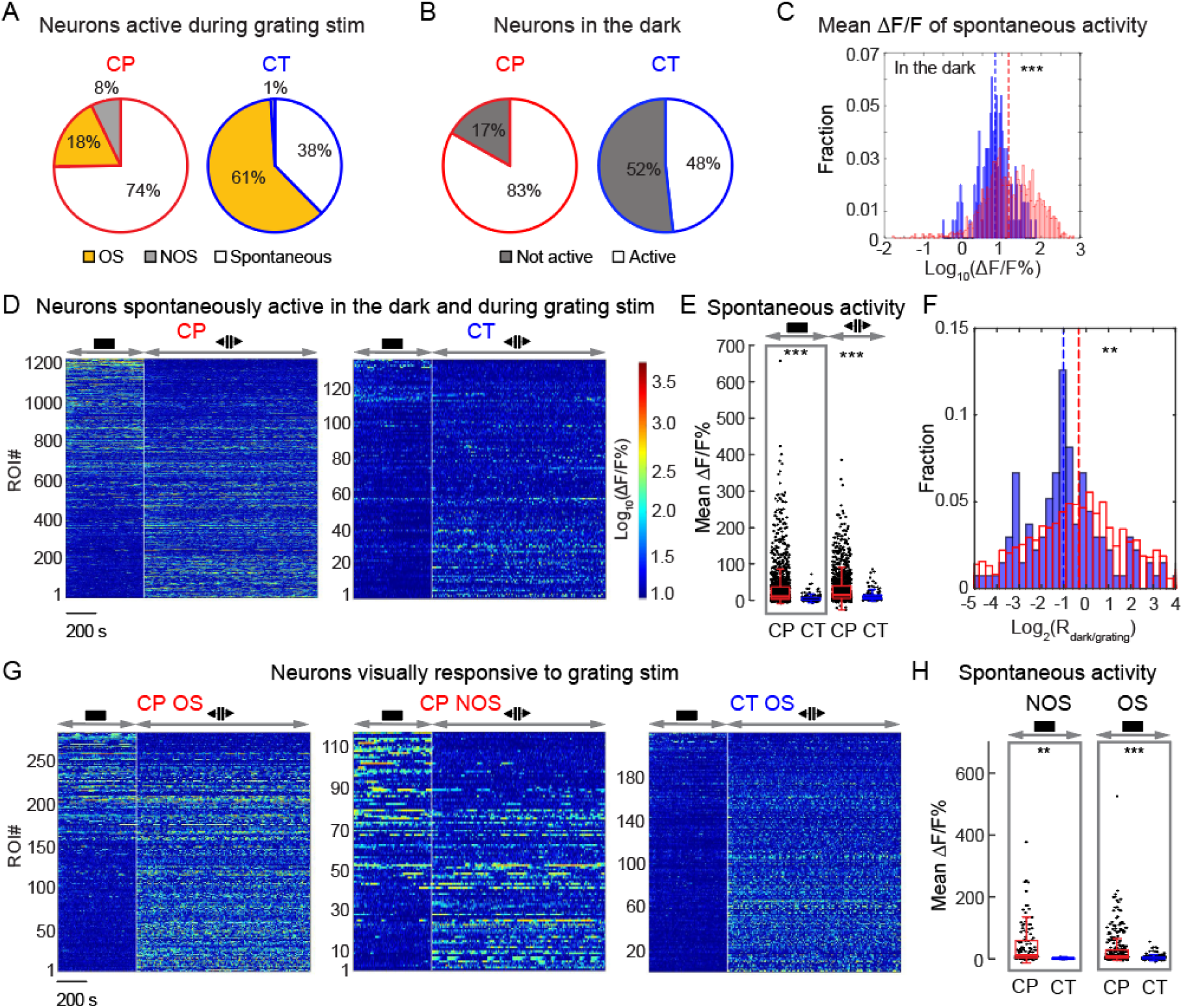
V1 L6 CP neurons exhibit pervasive spontaneous activity both in the dark and under visual stimulation. (A) Percentages of active CP and CT neurons with non-visually evoked, visually-evoked OS, and visually-evoked NOS responses, respectively, during drifting grating stimuli. CT: 1282 neurons from 10 mice; CP: 1876 neurons from 17 mice. Active neurons if its 99% ΔF/F value is larger than 50%; Visually evoked neurons if its activity during at least one visual stimulus was significantly higher than its activity during the inter-stimulus period by ANOVA test (p < 0.01). (B) Percentages of spontaneously active GCaMP6^+^ neurons in the dark. CT: 359 neurons from 5 mice; CP: 1625 neurons from 11 mice. (C) Histogram distributions of the mean ΔF/F% from the spontaneously active CP (red) and CT (blue) neurons in the dark. CT: 173 neurons from 5 mice; CP: 1351 neurons from 11 mice. (D) Raster plots of calcium transients associated with non-visually-evoked spontaneous activity of CP and CT neurons in the dark and under immediate subsequent drifting grating stimulation. ΔF/F% in log10 scale. Neurons were sorted according to the ratio of averaged ΔF/Fs in the dark and under drifting grating stimulation. CP: 1224 from 11 mice; CT: 135 neurons from 5 mice. (E) Scattered plots of the mean ΔF/F% for neurons in (D). (F) Histogram distributions of the ratios of mean ΔF/F% in the dark and under grating stimuli (R_dark/grating_ in log2 scale) for the CP (red) and CT (blue) neurons in (D). (G) Raster plots of calcium transients (ΔF/F% in log10 scale) and (H) mean ΔF/F% associated with spontaneous activity in the dark for CP (red) and CT (blue) neurons with visually evoked activity. CP: 284 OS and 117 NOS neurons from 11 mice; CT: 218 OS and 6 NOS neurons from 5 mice. Wilcoxon rank sum test: ***p<0.001 and **p<0.01.

Next we imaged the same neurons with the mice first kept in the dark and then presented with drifting grating stimuli. We identified CP and CT neurons that did not have visually evoked activity but only were spontaneously active both in the dark and during grating stimuli for further analysis (1224 CP neurons, 135 CT neurons, **Figure 5D-F**). Similar to above (**Figure 5C**), the spontaneous activity of CP neurons in the dark and during grating stimulus presentation had larger transients than the CT population (mean ΔF/F %, median values in the dark: CP 10.4 vs CT 3.6, p < 0.001; median values during grating: CP: 14.8 vs CT: 7.4, p < 0.001, Wilcoxon rank sum test; **Figure 5E**). Interestingly, even though these neurons did not exhibit visually evoked activity, the spontaneous calcium transients of a large proportion of CP neurons (68%) were strongly modulated by the presence of grating stimuli (**Figure 5D**): these CP neurons either increased (29%) or decreased (40%) their mean ΔF/F by at least 2-fold when the animal transitioned from being in the dark to being presented with grating stimuli. Although similar percentage (66%) of CT neurons had a ≥2× change of their spontaneous activity level during the transition from dark to grating sessions, more of them exhibited increasing mean ΔF/F% when grating stimuli were presented (51% with ≥2× activity gain vs. 15% with ≥2× with activity reduction, **Figure 5F**). By definition, such spontaneous activity was not directly evoked by visual stimuli. The observed modulations by the stimulation condition, however, suggest that they may reflect changes in the internal states that resulted from changes of the animal’s sensory perception.

We also found prominent differences in spontaneous activity of CP and CT neurons that showed visually evoked activity to grating stimuli (**Figure 5G**). CT neurons exhibiting visually-evoked responses had little spontaneous activity in the dark. In contrast, both OS and NOS CP neurons had much stronger spontaneous calcium transients in the dark (mean ΔF/F %, OS neurons: CP 7.1, 284 cells from 11 mice, CT 2.9, 218 cells from 5 mice, p < 0.001; NOS neurons: CP 10.4, 117 cells from 11 mice, CT 0.7, 6 cells from 5 mice, p < 0.01; Wilcoxon rank sum test, **Figure 5H**), indicating pervasive spontaneous activity as a distinct feature of L6 CP neurons. Together with the CP neurons that encoded visual information, through the cross-callosal horizontal network observed in our anatomical experiments, CP neurons thus convey both visual and non-visual information to multiple cortical regions across two hemispheres.

### Activity correlation of CP and CT neurons with arousal level

Having discovered that L6 neurons exhibit strong spontaneous activity, we further investigated how their activity was correlated with the arousal level of the animal. Given that pupil diameter is a well-established measure of arousal level, with enlarged pupil correlating with heightened alertness^35^, we combined simultaneous pupillometry recording with two-photon *in vivo* imaging of L6 neurons in the awake mouse (**Figure 6A**), where the mouse was first kept in the dark and then presented with drifting grating stimuli (**Figure 6B**).

**Figure 6.**
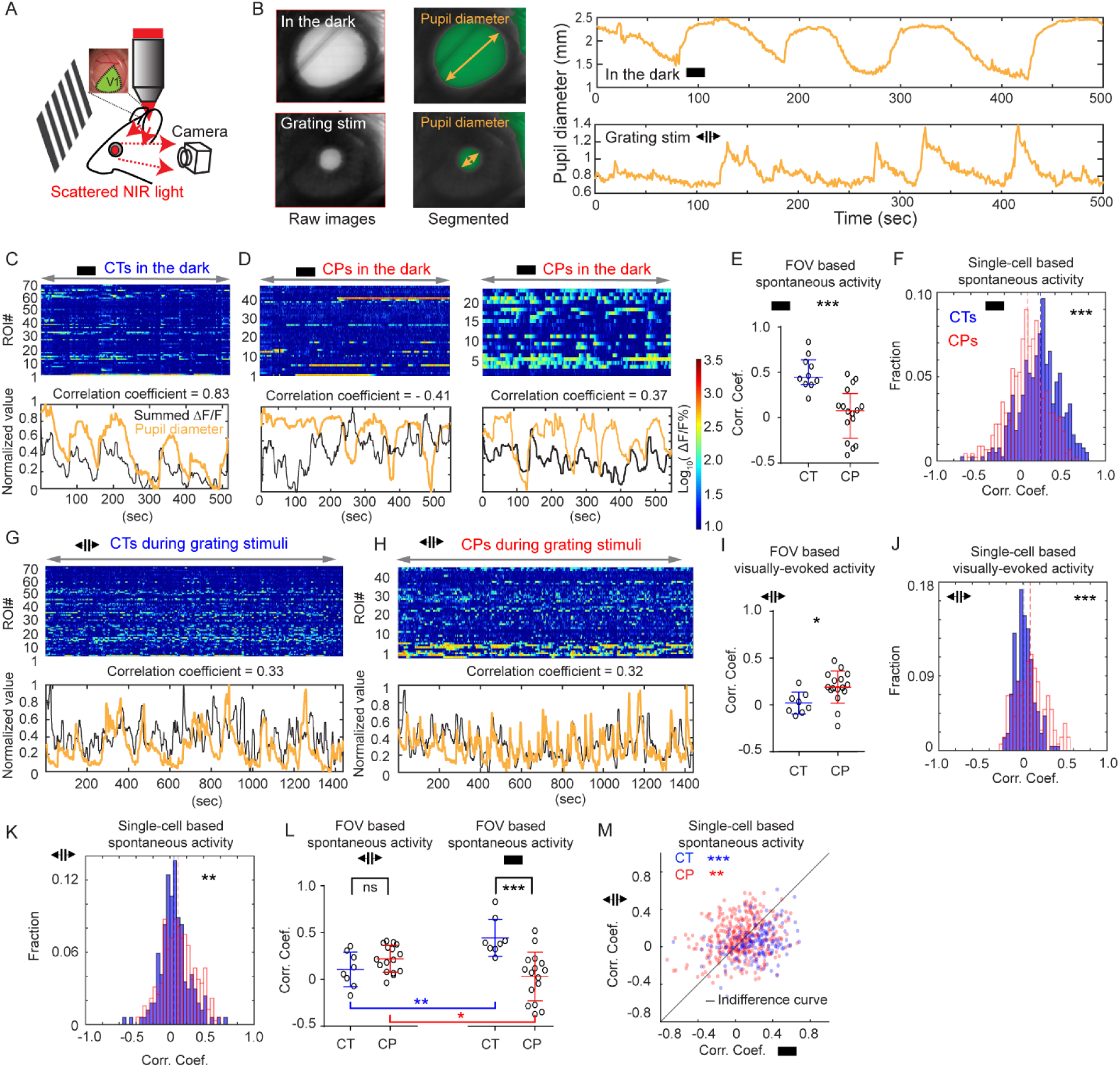
Activity of L6 CP and CT neurons shows distinct patterns of correlation with arousal level. (A) Arousal level was evaluated by pupillometry, where pupil was imaged using the near IR (NIR) two-photon excitation light scattered through the brain and emitted from the eye simultaneously with *in vivo* calcium imaging. (B) Example pupillometry results showing the pupil in the dark (upper panels) and during grating stimuli (lower panels). From left to right: raw image, segmented image, and temporal dynamics of pupil diameter. (C, D) Upper panels: Spontaneous Ca^2+^ response (ΔF/F %) of all active CT and CP cells in (C) one (for CT) and (D) two (for CP) example FOVs with the animals in the dark. Lower panels: Normalized pupil diameter (orange) and summed ΔF/F of all neurons in the FOV (black). Correlation coefficients were calculated between pupil diameter and summed ΔF/F. (E) FOV-based comparison of correlation coefficients between pupil diameter and summed spontaneous activity of each FOV of CT (10 FOVs from 5 mice) and CP (16 FOVs from 5 mice) neurons in the dark. p < 0.001, Mann-Whitney U test. (F) Histogram distribution comparison of correlation coefficients between pupil diameter and spontaneous activity of individual CT and CP cells (384 CT neurons from 5 mice and 456 CP neurons from 5 mice) in the dark. p < 0.001, Mann-Whitney U test. (G, H) Upper panels: Ca^2+^ response (ΔF/F %) of all active cells (including visually evoked and spontaneous activity) in example (G) CT and (H) CP FOVs with the animals under grating stimulation. Lower panels: Normalized pupil diameter (orange) and summed ΔF/F of all neurons in the FOV (black). Correlation coefficients were calculated between pupil diameter and summed ΔF/F. (I) FOV-based comparison of correlation coefficients between pupil diameter and summed visually evoked activity of each FOV of CT (8 FOVs from 5 mice) and CP (16 FOVs from 5 mice) neurons during grating stimulation of the animal. p < 0.01, Mann-Whitney U test. (J) Histogram distribution comparison of correlation coefficients between pupil diameter and visually evoked activity of visually-evoked cells during the grating stimulation (215 CT neurons from 5 mice and 136 CP neurons from 5 mice), p < 0.001, Mann-Whitney U test. (K) Histogram distribution comparison of correlation coefficients between pupil diameter and spontaneous activity of non-visually-evoked cells during the grating stimulation (169 CT neurons from 5 mice and 320 CP neurons from 5 mice). p < 0.01, Mann-Whitney U test. (L) FOV-based comparison of correlation coefficients between pupil diameter and summed spontaneous activity of each FOV of CT (8 FOVs from 5 mice) and CP (16 FOVs from 5 mice) neurons (left panel) during grating stimulation and (right panel) in the dark. ***p < 0.001, **p < 0.01, *p < 0.05, ns p > 0.05, Mann-Whitney U test for CT and CP comparison under the same stimulus condition, Wilcoxon signed-rank test for comparing the same CT or CP FOVs under two stimulus conditions. (M) Single-cell based comparison of correlation coefficients between pupil diameter and spontaneous activity (y-axis) during grating stimulation or (x-axis) in the dark of CT (in blue) or CP (in red) neurons that were spontaneous active in the dark and during grating stimulation (169 CT neurons from 5 mice and 320 CP neurons from 5 mice). *** p < 0.001, ** p < 0.01, Wilcoxon signed-rank test.

We first studied how pupil diameter was correlated with spontaneous activity in the dark (**Figure 6C-F**). At the population level, we calculated the Pearson correlation coefficients between pupil diameter and the summed ΔF/F of all active CT or CP cells in the same FOV within individual imaging sessions (Example FOVs in **Figure 6C** and **6D**). We found that, in the dark, the population activity of CT neurons in each FOV was consistently positively correlated with pupil diameter. In contrast, the population activity of CP neurons displayed a wide variation in their correlation with pupil diameter, with imaging sessions where the FOV activity exhibited positive, negative, or a lack of correlation with arousal. As a result, CP neuron FOVs had significantly lower correlation coefficients than CT (median of correlation coefficients: 0.44 for CT vs 0.07 for CP, 10 FOVs from 5 CT mice, 16 FOVs from 5 CP mice, p < 0.001, Mann-Whitney U test, **Figure 6E**). We observed the same trend on single-cell level and found the spontaneous activity of individual CT neurons to be more likely to exhibit positive correlation with arousal than CP neurons (medians: 0.25 for CT vs 0.08 for CP, 384 CT neurons from 5 mice and 456 CP neurons from 5 mice, p < 0.001, Mann-Whitney U test, **Figure 6F**).

We then measured CT and CP neuronal activity during grating stimulation (**Figure 6G-K,** example FOVs in **Figure 6G** and **6H**). Because CT and CP neurons may exhibit either visually evoked or spontaneous activity during grating stimulation, we divided neurons into two populations based on whether they showed visually evoked activity and separately studied their activity correlation with arousal. For neurons with visually evoked activity, on both population and single-cell levels (**Figure 6I** and **6J**), CP neurons were more likely to be positively correlated with arousal, whereas CT neurons’ correlation with arousal averaged around zero (**Figure 6I**, median correlation coefficients of visually evoked activity at FOV level: 0.18 for CP vs −0.001 for CT, 16 FOV from 5 mice for CP, 8 FOVs from 5 mice for CT, p < 0.05; **Figure 6J**, median correlation coefficients of visually evoked activity at single cell level: 0.08 for CP vs 0.009 for CT, n = 136 for CP from 5 mice, n = 215 from CT from 5 mice, p < 0.001; Mann-Whitney U test). For spontaneous activity during grating stimulation, although on single-cell level there was a significantly higher fraction of CP neurons showing positive correlation with arousal than CT (**Figure 6K**, median correlation coefficients for spontaneous activity: 0.09 for CP vs 0.05 for CT, n = 320 for CP from 5 mice, n = 169 from CT from 5 mice, p < 0.01, Mann-Whitney U test), on the population FOV level, difference between CT and CP was not significant (left panel of **Figure 6L**, median correlation coefficients: 0.21 for CP vs 0.09 for CT, 16 FOV from 5 mice for CP, 8 FOVs from 5 mice for CT, p = 0.14, Mann-Whitney U test).

Comparing with the spontaneous activity of the same neurons in the dark (right panel of **Figure 6L**, FOV comparison, median correlation coefficients of CPs: 0.086 in dark vs 0.21 during grating, p < 0.05; for CTs: 0.39 in dark vs 0.09 during grating, p < 0.01; 16 FOV from 5 mice for CP, 8 FOVs from 5 mice for CT, Wilcoxon signed-rank test; **Figure 6M**, single-cell comparisons, median correlation coefficients of CPs: 0.078 in dark vs 0.094 during grating, p < 0.01; for CTs: 0.25 in dark vs 0.046 during grating, p < 0.001; 320 CP neurons from 5 mice and 169 CT neurons from 5 mice, Wilcoxon signed-rank test), we found intriguing differences between CT and CP neurons. For spontaneous activity of CT population, its correlation with arousal substantially decreased when the mouse transitioned from being in the dark to being under grating stimulation. For CP neurons, however, its spontaneous activity correlation with arousal followed the opposite trend and went from being highly heterogeneous with both negative and positive correlation to being mostly positive correlation. Together, our investigations on the spontaneous activity characteristics of L6 neurons indicated that CP neurons possessed pervasive spontaneous activity during both the absence and the presence of strong visual input, with activity patterns and correlation with arousal distinct from those of CT neurons. Together with the anatomical results presented earlier, they point to new and yet-to-explored functions of these cross-callosal neurons.

## Discussion

Despite a growing recognition of the prominent roles that the corpus callosum plays in the processing of sensory information, the identity and property of CP neurons are not well understood. Traditionally considered to be L2/3 or L5 neurons projecting homotopically to the contralateral cortex, CP neurons were thought to contribute to the processing of sensory information encoding bilateral stimuli. In this work, using a designer variants of recombinant adeno-associated virus, rAAV2-retro, we gained selective genetic access to CP neurons projecting to mouse V1 and investigated their connectivity pattern and functional properties.

We found that in contrast to L2/3 and L5 CP neurons at V1/V2 border (corresponding to the binocular visual field) that contribute to binocularity, within monocular V1, the main CP neurons mediating cross-callosal communication were a population of L6 neurons that were distinct from the more well-known thalamus-projecting L6 CT neurons. Immunostaining with GABA confirmed the excitatory nature of the CP neurons, consistent with previous studies in cat and rat^36^. Although we found the ratio of CP to CT neurons to be one to nine, the proportion of CP neurons was almost certainly underestimated due to incomplete retrograde labeling. In the context of L6 neurons in general, the CP L6 neurons identified here belong to the corticocortical (CC) population that provide only corticocortical but not corticothalamic projections. In the rat primary somatosensory cortex, an equal proportion of CT and CC neurons were found, implicating L6 as a substantial contributor of corticocortical projections^25^. Our results further indicate that the CP subpopulation of the L6 CC neurons subserve cross-callosal communications for monocular V1.

Contralateral V1 is not the only source of cross-callosal inputs to V1. We found that L6 CP neurons from the secondary visual cortices and L5 CP neurons from primary auditory cortex projected across corpus callosum into V1. Using the same viral strategy, we also discovered that V1 L6 CP neurons projected back to these sensory areas. Therefore, these cross-callosal projections are not homotopic but spanning multiple cortical regions. We found the main cross-callosal axonal projections to localize in the infragranular layers of V1 and that these projections formed monosynaptic connection with the ipsilateral CP neurons, indicating direct reciprocal connectivity between CP neurons of the two hemispheres. Together, our results revealed the existence of an extensive cross-callosal reciprocal network mediated by L6 neurons in the visual cortical areas. The involvement of higher visual cortices in this pathway suggests the presence of higher-level visual representations, which may provide contextual information to early processing of visual information in V1. The inclusion of AuC within this reciprocal network leads us to speculate that this network is also involved in cross-modal interactions across the hemispheres^37^. Indeed, a recent work reported that visual stimuli can directly evoke activity of L6 neurons in auditory cortex^38^, which can be partly attributed to the cross-callosal inputs from L6 CP neurons in visual cortices. Similarly, by conveying auditory information from the contralateral hemisphere, the transcallosal network described here can also contribute to the contextual modulation of V1 by auditory signals^39, 40^.

The retrogradely transported rAAV2-retro also allowed us to express genetically encoded calcium indicators in V1 CP neurons and study their visually evoked as well as spontaneous activity in awake mice *in vivo*. With drifting grating stimuli, we found both CP and CT neurons with visually evoked responses with some subtle but significant differences in their response properties: whereas almost all visually-driven CT neurons were OS, about 70% of CP neurons were; CT neurons were more direction selective than CP neurons; the fraction of visually responsive CT neurons that had well-defined receptive fields was also about double that of the CP neurons (60% vs. 30%); both populations included sharply orientation-tuned cells, although CP also included more broadly tuned neurons. Despite these differences, our experiments indicate that, by encoding visual features such as orientation, direction, and receptive field, L6 CP neurons convey information on the monocular visual field across corpus callosum into the contralateral hemisphere.

More distinct are the spontaneous activity patterns of CP and CT neurons. Investigating how their activity related to arousal level, we found that in the dark, spontaneous population activity of *NTSR1*-positive CT neurons was strongly positively correlated with the arousal level of the mouse. Given that these *NTSR1*-positive CT neurons are known to be directly depolarized and potently modulated by acetylcholine^41^, in the absence of visual inputs the spontaneous activity that we observed in the dark was likely strongly modulated and/or driven by the cholinergic inputs into V1. Upon drifting grating stimulation, CT population activity correlation with arousal became significantly less than in the dark, possibly because the strong visual inputs that they received from local V1 neurons reduced the influence of and thus the correlation with cholinergic activity. Together with recent results where CT neurons were found to control the gain of local visually evoked cortical activity^22^, our observation also suggests a functional pathway for arousal level to modulate V1 activity.

Compared with CT neurons, the activity patterns for CP population were more dominated by spontaneous activity. Both during drifting grating stimuli and with the mice kept in the dark, CP neurons were much more likely than CT neurons to exhibit spontaneous activity (74% vs. 38% during grating stimuli, 83% vs. 48% in the dark) with larger calcium transient magnitudes. The heightened spontaneous activity of CP population may be explained by its presynaptic input pattern. Whereas CT and CP populations both receive local V1 inputs, we found CP neurons to receive more long-range inputs from higher cortical areas of the ipsilateral hemisphere, which are likely feedback in nature and known to target deeper layers^42, 43, 44^. They also received cross-callosal cortical inputs via the extensive CP neuron network described above. Given the beliefs that spontaneous activity is primarily driven by corticocortical connections^45, 46, 47^ and projections from higher cortical regions initiate spontaneous patterns in deep layers of primary sensory cortex^48^, these additional sources of cortical inputs for CP neurons may account for their heightened spontaneous firing.

These long-range cortical inputs may also cause CP neurons to have more diverse activity correlations with brain states. Unlike CT neurons, whose spontaneous population activity in the dark was strongly positively correlated with arousal, the spontaneous activity of the CP population in the dark may be negatively, positively, or un-correlated with the arousal level, indicating more complex (e.g., multisensory, higher cognitive) origins of forces that drive and modulate their spontaneous firing^49^. During grating stimuli, the population activity of CP neurons becomes more positively correlated with arousal than in the dark, suggesting a switch of cortical dynamics and brain state when the animal was exposed to strong sensory stimulation.

For CT and CP neurons with spontaneous activity both in the dark and during grating stimuli, we found that their activity, albeit not temporally synchronized with stimulus onsets and thus not directly visually evoked, was modulated by the presence of the grating stimuli. These parallel the observations made in alert macaque V1, where L6 were found to be the dominant spontaneously active layer both in the dark and in the light, with the change of illumination condition modulating their firing rates^50, 51, 52^. Such changes of spontaneous firing rate may encode the presence, timing, or luminance of a stimulus, as well as reflect the associated shifts of the animal’s cortical and behavioral states.

Previous studies indicated that spontaneous events often originate from infragranular layers then spread upwards into superficial layers^48, 53^. In addition to leading to spontaneous activity in local V1 circuit and ipsilateral cortical regions, CP neurons also convey information to contralateral hemisphere through the corpus callosum, thus can influence cortical dynamics more globally. The exact functional roles of the spontaneous activity observed by us are unknown. Nevertheless, with their visually driven as well as spontaneous activity and given their extensive reciprocal network spanning multiple cortical areas in two hemispheres, L6 CP neurons are the ideal candidates in broadcasting both visual and nonvisual information globally, thus regulating and coordinating brain-wide activity events from sensory perception to memory replay ^54, 55, 56^.

## Methods

All experimental protocols were conducted according to the National Institutes of Health guidelines for animal research and approved by the Institutional Animal Care and Use Committee at Janelia Research Campus, Howard Hughes Medical Institute.

### Experimental Model and Subject Details

The following mouse lines were used: Wild-type C57BL/6J (Jackson Laboratory); NTSR1-Cre (strain B6.FVB(Cg)-Tg(NTSR1-cre)GN220Gsat/Mmcd, stock number 030648-UCD); Scnn1a-Tg3-Cre mice (Jax no. 009613); Rbp4-Cre mice (MMRRC no. 031125-UCD); Gad2-IRES-Cre (Jax no. 010802); tdTomato reporter line (Ai14, Jax. 007908); nuclear tdTomato reporter line (R26 LSL H2B mCherry 1H3 line, Jax no. 023139); Ai3 mice (JAX Stock No: 007903). Mice of both sexes (older than P60) were used. Sample sizes (number of mice, cells and/or field-of-view, FOVs) for each experiment are stated in main text. AAV viruses were obtained from Virus Services of Janelia Research Campus, HHMI.

### Virus injection for histology

Virus injection and cranial window implantation procedures have been described previously ^57^. Briefly, mice were anaesthetized with isoflurane (1–2% by volume in O_2_) and given the analgesic buprenorphine (SC, 0.3 mg per kg of body weight). Virus injection was performed using a glass pipette (Drummond Scientific Company) beveled at 30° with a 15 to 20-μm opening and back-filled with mineral oil. A fitted plunger controlled by a hydraulic manipulator (Narashige, MO10) was inserted into the pipette and used to load and inject the viral solution.

For the injection of virus for histological examination, a burr hole was made (∼200 µm diameter) over the injection site. 30 nl virus-containing solution (rAAV2-retro.CAG.GFP, 1×10^13^ infectious units per ml) was injected 0.6 mm below pia at two injection sites for each brain region. The injection coordinates for each brain region are: (i). V1: midline: 2.5 mm, Bregma: −3.4 mm and −4.0 mm; (ii). V2M: midline: 1.25 mm, Bregma: −3.4 mm and −4.0 mm; (iii). V2L: midline 3.5 mm, Bregma: −3.4 mm and −4.0 mm; (iv). Auditory cortex: midline 4.0 mm, Bregma: −3.4 mm and −4.0 mm.

For rabies tracing experiment, 1:1 mixture of AAV2/1.CAG.FLEX.BFP.T2A.TVA (5×10^13^ infectious units per ml) and AAV2/1.CAG.FLEX.BFP.T2A.G (7.2×10^12^ infectious units per ml) were injected into left V1 at the coordinates described above (30 nl at 0.6 mm below pia). For the labeling of CP neurons, rAAV2-retro.syn.Cre (1×10^13^ infectious units per ml) was first injected into right V1 at the same coordinates, before rabies vectors were injected. Three weeks later, ΔRV.mCherry (3.4×10^8^ infectious units per ml) was injected into the same injection sites in left V1 (30 nl, 0.6 mm below pia).

### Viral injection and cranial window implantation for *in vivo* imaging

For the labeling of CT and CP cells for calcium imaging, a 3.5-mm diameter craniotomy was first made over left V1 of NTSR1-Cre and wildtype mice, respectively. Then 30 nl of virus-containing solution (AAV2/1.syn.FLEX.GCaMP6s, 1× 10^13^ infectious units per ml) was injected 0.6 mm below pia into left V1 at four injection sites at the intersection points of the two left-right lines at Bregma −3.4 mm and −4.0 mm, and two anterior-posterior lines at 2.2 mm and 2.6 mm from the midline. For the labeling of CP neurons, rAAV2-retro.syn.Cre (1× 10^13^ infectious units per ml) was first injected into the contralateral V1 at the same coordinates, before craniotomy was performed. For dual-color imaging experiments, a 1:1 mixture of AAV2/1.CAG.FLEX.jRGECO1a and AAV2/1.CAG.FRT.GCaMP6s was injected into left V1 of NTSR1-Cre mice, and rAAV2-retro.syn.FLPo was injected into the right V1 of the same animal. After the pipette was pulled out of the brain, a glass window made of a single coverslip (Fisher Scientific, no. 1.5) was embedded in the craniotomy and sealed in place with dental acrylic. A titanium headpost was then attached to the skull with cyanoacrylate glue and dental acrylic.

### Visual stimulation

Visual stimuli were presented by back projection on a screen made of Teflon film using a custom-modified DLP projector. The screen was positioned 17 cm from the right eye, covering 75° × 75° degrees of visual space and oriented at ∼40° to the long body axis of the animal. The projector provided equilength and linear frames at 360 Hz (designed by A. Leonardo, Janelia Research Campus, and Lightspeed Design, model WXGA-360). Its lamp housing was replaced by a holder for liquid light guide, through which visible light (450–495 nm) generated by a LED light source (SugarCUBE) was delivered to a screen made of polytetrafluoroethylene. The maximal luminance measured at the location of animal eyes was 437 nW/mm^2^. Visual stimuli were generated using custom-written codes. During visual stimulation, the luminance level was kept constant. To measure orientation-tuning, full-field square gratings were presented in 12 directions in a pseudorandom sequence for 12 s each, during which time each stimulus was static for the first and last 3 s and moving during the middle 6 s. Gratings had 100% contrast, 0.07 cycles per degree, and drifted at 26 degrees per second (i.e., a temporal frequency of ∼2 Hz). Each oriented grating was presented for a total of ten trials.

### Pupil tracking

An infrared-sensitive CCD camera controlled by a custom written interface with LabVIEW® collected images of the pupil illuminated by two-photon excitation light scattered into the eye during brain imaging (Figure 7A) at 10-Hz frame rate. The pupil was segmented by custom-written MATLAB codes and pupil diameter values interpolated to find the diameter of the pupil corresponding to each two-photon image.

### Two-photon imaging

All imaging experiments were carried out on head-fixed, awake mice, except data in Figure S4, for which mice were anesthetized for receptive field mapping. To habituate the mice to experimental handling, each mouse was head-fixed onto the sample stage with its body restrained under a half-cylindrical cover, which reduced struggling and prevented substantial body movements such as running. The habituation procedure was started one week after surgery, repeated 3–4 times for each animal, and each time for 15–60 min. Imaging was performed with two-photon fluorescence microscope 3–4 weeks after virus injection. Each experimental session lasted 45 minutes to 2 hours. Multiple sections (imaging planes) may be imaged within the same mouse. GCaMP6s was excited at 940 nm with a femtosecond laser (InSight Deepsee, Spectra-Physics) that was focused by either a Nikon 16×, 0.8 NA or an Olympus 25×, 1.05 NA objective. Emitted fluorescence photons reflected off a dichroic long-pass beamsplitter (FF665-Di02-25×36; Semrock) and were detected by a photomultiplier tube (H7422PA-40, Hamamatsu). jRGECO was excited at 1100 nm with the same laser source. For simultaneous imaging of GCaMP6s and jRGECO, 1030 nm was used for excitation.

Images of CP or CT neurons were acquired from 550 to 650 μm below pia. Laser power measured post objective varied between 67 mW and 329 mW (n = 38 imaging sessions from 17 mice). Typical time for mapping the orientation selectivity of a single image section was ∼25 min, during which no photobleaching or photodamage was observed. Typical images had 256 × 256 pixels, at 1.2–2.2 μm per pixel and 2-3 Hz frame rate.

### Histology

The survival time before histological evaluation for mice injected with tracers (FG) and AAVs was one week and three weeks, respectively. For mapping of presynaptic neurons, Cre-dependent AAVs encoding rabies glycoprotein (G) and the avian virus receptor (TVA) were injected into mice with target cells expressing Cre. Three weeks later, modified rabies virus (ΔRV) encoding mCherry was injected into the same mouse, resulting in the targeted infection of the previously labeled neurons, and subsequent trans-synaptic spread and expression of mCherry. The brain was fixed after 8 days post rabies virus injection. For histological examination, mice were deeply anaesthetized with isoflurane and transcardially perfused with PBS and then 4% paraformaldehyde (w/v). Brains were removed and post-fixed overnight in 4% paraformaldehyde. Fixed whole brains were embedded in 4% agar and sliced with vibrating microtome (V1200S, Leica) at the thickness of 100 μm for direct observation or 40 μm for immunostaining. We performed immunostaining by application of primary antibodies (overnight): chicken-anti-GFP (Aves, 1:200) for GCaMP6s, or anti-GABA (Abcam, 1:200) to identify interneurons. After three washes for 5 min each in PBS, secondary antibodies were applied along with 0.1% Triton X-100 for 1 hr. For secondary antibodies, we used Alexa Fluro 488-conjugated donkey anti-chicken (Invitrogen, 1:500) or Alexa Fluro 594-conjugated donkey anti-rabbit (Invitrogen, 1:500). All brain slices were mounted in Vector Shield mounting solution. Coronal images were acquired via a stereomicroscope at low zoom (2-4×), at high zoom with Zeiss ApoTome.2 (20×/0.8NA, optical section step of 0.5 μm), or on a confocal microscope (Zeiss LSM 800, 63×/1.4 NA oil immersion, optical section step of 0.5 μm). For rabies tracing, cells were manually counted on individual coronal slices with the brain region determined using a standard mouse brain atlas ^58^. We counted 282±56, 409±76, and 242±42 mCherry (+) cells (mean ± s.e.m.) when CT, CP, and L4 pyramidal cells were starter cells, respectively.

### Analysis of Two-Photon Imaging data

Imaging data were processed with custom programs written in MATLAB (Mathworks®) and Fiji ^59^. Images were registered with an iterative cross-correlation-based registration algorithm ^57^. Cortical neurons were outlined by hand as regions of interest (ROIs). The averaged fluorescent signal within the ROI was used to calculate calcium transients. For each ROI, we used the mode from the fluorescence intensity histogram as the baseline fluorescence F_0_, and calculate its calcium transient as ΔF/F (%) = (F−F_0_)/F_0_ × 100. The final calcium transient to each visual stimulus was the average of ten trials.

We calculated the mean of the ΔF/F values that were 99% or above in the calcium transient distribution of each ROI during an imaging session. A neuron was considered active if this mean value of its corresponding ROI was above 50%. We compared this criterion with our previously used one (the maximum of mean ΔF/F during the presentation of visual stimuli was above 10%^57^) on data of CT neurons in the drifting grating sessions (1503 cells, 10 mice). We found very similar percentages of active neurons (85.3% versus 83.6%, new versus old criterion), suggesting that the two criteria were equivalent in identifying active neurons. Since the new criterion allowed the evaluation of spontaneous activity, we used this criterion throughout our analysis. Of 2066 CP cells outlined from 17 mice, 1876 (90.8%) were active. A neuron was considered as visually-evoked if its activity during at least one visual stimulus was significantly higher than their activity during the inter-stimulus period by paired t test (P < 0.01) ^31^. Under this criterion, the percentages of visually-evoked neurons among all active neurons were: CT 62%, 796/1282, 10 mice; CP 26%, 483/1876, 17 mice.

### Analysis of Orientation Selectivity of individual neurons

The orientation selectivity index (OSI), directional selectivity index (DSI), global OSI, global DSI, and tuning full width at half maxima (FWHM) was defined based on previous publications^57, 60, 61^. Briefly, the response R of each ROI to a visual stimulus was defined as the average ΔF/F across the 6-s window of drifting grating presentation. For ROIs with significantly different responses across the drifting directions (one-way ANOVA, P < 0.05), we fit their normalized response tuning curves to grating drifting angle θ with a bimodal Gaussian function. The tuning width for the preferred orientation is calculated as the full width at half maximum (FWHM) of the Gaussian function. OSI was computed as (*R*_pref_−*R*_ortho_)/(*R*_pref_+*R*_ortho_), with *R*_pref_ and *R*_ortho_ being the responses at the preferred and orthogonal orientations, respectively. With this index, perfect orientation selectivity would give OSI=1; an equal response to all orientations would have OSI=0. DSI was defined as (*R*_pref_−*R*_ortho_)/(*R*_pref_+*R*_ortho_), where *R*_pref_ and *R*_ortho_ are the responses at the preferred motion direction and its opposite, respectively. Global OSI was calculated as the magnitude of the vector average divided by the sum of all responses: gOSI = lΣ_k_R(θ_k_)e^i2θ_k_^l/Σ_k_R(θ_k_), where R(θ) is the measured response at orientation θ, and global DSI was defined as 1Σ_k_R(θ_k_)e^iθ_k_^l/Σ_k_R(θ_k_).

### Statistical Tests

Standard functions and custom-written scripts in MATLAB were used to perform all analysis. The data were tested for normal distribution. Parametric tests were used for normally distributed data and non-parametric tests were applied to all other data. Bar graphs and mean ± SEM were used to describe the data with normal distribution, while boxplots and median ± IQR were used to describe the non-normally distributed data. Boxplots represent median and 25^th^ - 75^th^ percentiles and their whiskers shown in Tukey style (plus or minus 1.5 times IQR). A nonparametric test (Wilcoxon signed rank test) was used to examine paired data in Figure 1F, Figure 7E and 7F. Direct non-paired comparisons between two groups were made using Wilcoxon rank sum test for non-normally distributed data (Figure 5I-5O, 6C, 6E, 6F, 6H). The statistical significance was defined as ∗p < 0.05, ∗∗p < 0.01, ∗∗∗p < 0.001, respectively. Experiments were not performed blind. Sample sizes were not predetermined by statistical methods, but were based on those commonly used in the field. Medians, IQR, means and SEM are reported throughout the text.

## Acknowledgements

We thank Shuo Chen, Jianhua Cang, and Hillel Adesnik for helpful suggestions on the manuscript, Amy Hu for histology, and Kim Ritola for virus preparation. Y.L., W.S., R.L., and N.J. were supported by Howard Hughes Medical Institute. M.C. was supported by National Natural Science Foundation of China (31700909). N.J. was supported by National Institutes of Health (U19NS107613).

## Author contributions

N.J., Y.L. and W.S. designed the research. W.S. established awake imaging. R.L. built the pupil tracking device. W.S. and R.L. contributed codes for analysis of data. Y.L. and W.J. performed imaging experiment. Y.L. injected and implanted mice and performed histology. Y.L. analyzed two-photon imaging data. M.C. conducted the whole-cell electrophysiological recordings supervised by W.S. N.J. supervised research. Y.L. and N.J. wrote the manuscript.

## Materials & Correspondence

All data are available from the Lead Contact, Na Ji, upon request.

