## Supplemental figures for "A distinct population of L6 neurons in mouse V1 mediate cross-callosal communication"

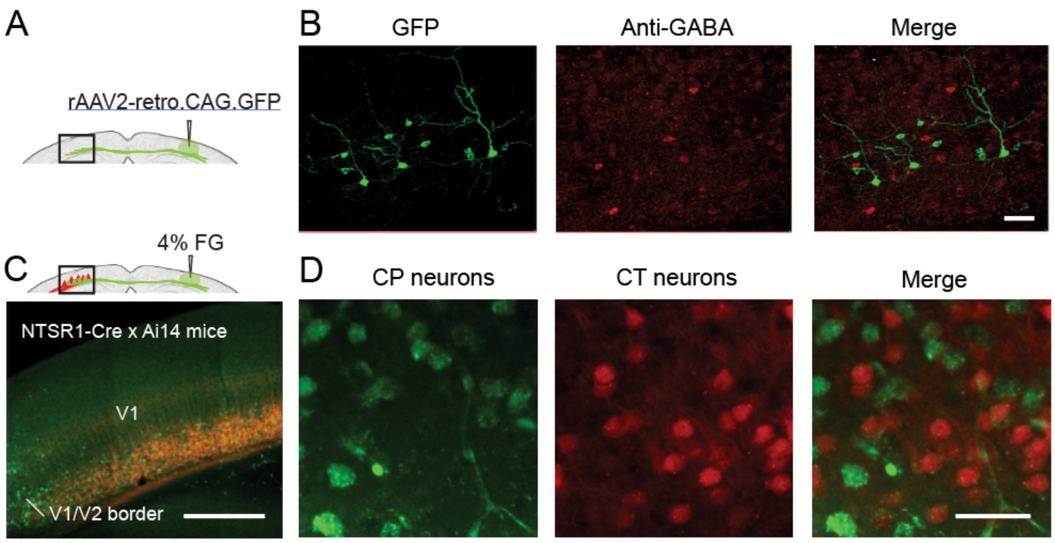


**Supplementary Figure 1. GABA immunostaining of L6 and retrograde tracing of callosal projection neurons in V1 by 4% FG**

(A) CP neurons were labeled by contralateral injection of rAAV2-retro.syn.GFP.

(B) Coronal brain sections immunostained with anti-GABA (in red) showed that L6 CP neurons were not immunoreactive to anti-GABA.

(C) 4% FG was injected into the right V1 of NTSR1-Cre × Ai14 mice where CT neurons expressed tdTomato. $FG^{+}$ (green) cells were found in L6 of the left V1.

(D) No colocalization was found between $FG^{+}$ cells (green) and CT neurons (red).

Scale bars: 50 μm in (B, D), 500 μm in (C).

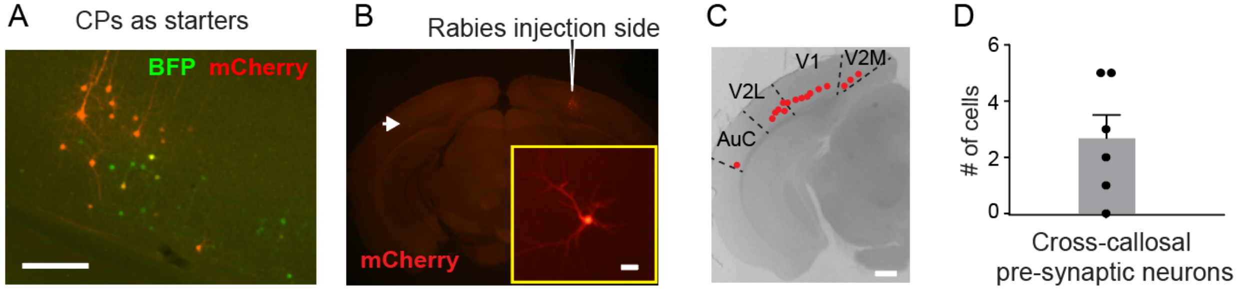


**Supplementary Figure 2. Monosynaptic connections between L6 CP neurons across hemispheres**

(A) Example coronal image of the rabies injection site with CP neurons as starter cells.

(B) Example coronal image showing a presynaptic cell (arrow) contralateral to the injection site. Inset: a magnified image of the cell.

(C) The locations of all presynaptic CP neurons (16 cells from 6 mice) contralateral to the injection site. Each red dot represents a cell.

(D) Numbers of presynaptic cells in the hemisphere contralateral to the injection site with CP as the starter cells. n = 6 mice.

Scale bars: 200 μm in (A), 500 μm in (B, C).

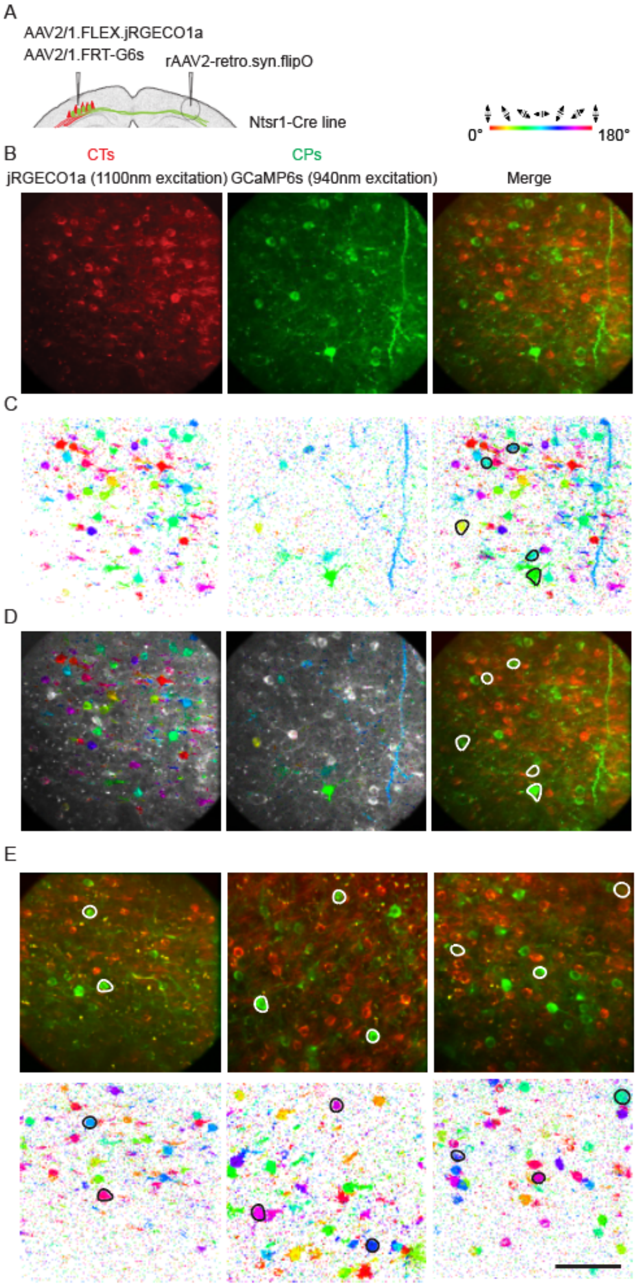


**Supplementary Figure 3. Simultaneous *in vivo* calcium imaging of CP and CT neurons and their tuning maps**

(A) Viral labeling strategy in NTSR1-Cre mice: CP neurons were labeled by contralateral injection of rAAV2-retro.syn.FLPo and ipsilateral injection of AAV2/1.FRT.GCaMP6s. CT neurons were labeled by ipsilateral injection of AAV2/1.syn.FLEX.jRGECO1a.

(B) Example *in vivo* two-photon excitation fluorescence images of L6 CT (red, jRGECO1a$^{+}$) and CP (green, GCaMP6s$^{+}$) neurons in the same FOV. Orientation selectivity test was performed separately on CP neurons (at 940 nm excitation for GCaMP6s) and on CT neurons (at 1100 nm excitation for jRGECO1a).

(C) Preferred orientation map of CP and CT neurons in (B) with each pixel color-coded by its preferred orientations. In the rightmost merged turning maps, OS CP neurons are outlined in black.

(D) Overlap of fluorescence images and turning maps. In the rightmost merged fluorescence images, OS CP neurons are outlined in white.

(E) Additional FOVs with merged channels and tuning maps.

Scale bar, 100 µm.

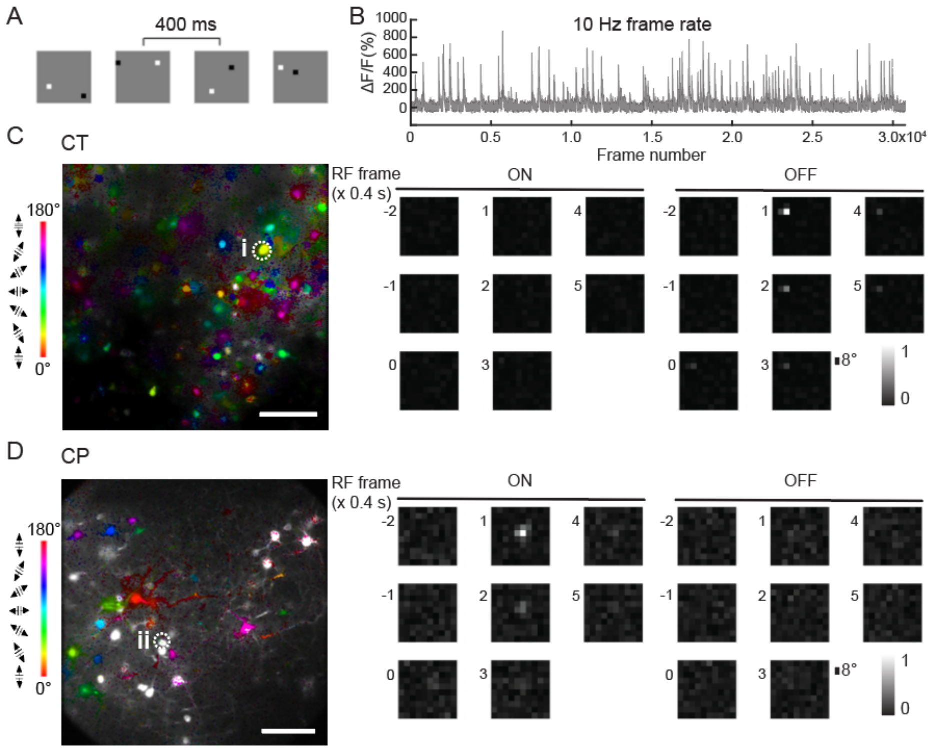


**Supplementary Figure 4. Receptive field mapping for CT and CP neurons**

(A) Receptive field (RF) mapping using sparse-noise stimuli of a pair of white ("ON" stimulus) and black ("OFF" stimulus) squares randomly distributed on a gray background presented 400 ms apart.

(B) Example calcium transient (ΔF/F%) trace from a CP neuron.

(C, D) Spatiotemporal RFs of an example L6 CT (C, ROI i) and CP (D, ROI ii) neurons, respectively. Left panels: orientation tuning maps; Right panels: "ON" and "OFF" RFs from 0.8 sec before to 2 sec after the start of visual stimulus ("0").

Scale bar: 100 μm in (C, D).
